# ProtoBind-Diff: A Structure-Free Diffusion Language Model for Protein Sequence-Conditioned Ligand Design

**DOI:** 10.1101/2025.06.16.659955

**Authors:** Lukia Mistryukova, Vladimir Manuilov, Konstantin Avchaciov, Peter O. Fedichev

**Affiliations:** GERO PTE. LTD., 133 Cecil Street 14-01 Keck Seng Tower, Singapore 069535

## Abstract

Designing small molecules that selectively bind to protein targets remains a central challenge in drug discovery. While recent generative models leverage 3D structural data to guide ligand generation, their applicability is limited by the sparsity and bias of structural resources. Here, we introduce ProtoBind-Diff, a structure-free masked diffusion model that conditions molecular generation directly on protein sequences via pre-trained language model embeddings. Trained on over one million active protein–ligand pairs from BindingDB, ProtoBind-Diff generates chemically valid, novel, and target-specific ligands without requiring structural supervision. In extensive benchmarking against structure-based models, ProtoBind-Diff performs competitively in docking and Boltz-1 evaluations and generalizes well to challenging targets, including those with limited training data. Despite never observing 3D information during training, its attention maps align with predicted binding residues, suggesting the model learns spatially meaningful interaction priors from sequence alone. This sequence-conditioned generation framework may unlock ligand discovery across the full proteome, including orphan, flexible, or rapidly emerging targets for which structural data are unavailable or unreliable.

## I. INTRODUCTION

The chemical space of drug-like molecules is estimated to exceed 10^60^ structures [1], making exhaustive exploration practically infeasible. Machine learning has emerged as a powerful tool to generate candidate compounds, guiding discovery beyond what traditional screening can reach. Generative AI models have been developed to address this challenge, leveraging various molecular representations, such as text strings, graphs, or 3D structures, and spanning a wide range of architectures, including transformers [2, 3], reinforcement learning agents [4], variational autoencoders (VAEs) and generative adversarial networks (GANs) [5, 6], and more recently, diffusion models [7, 8].

A promising yet challenging frontier is *protein-conditioned molecular generation*, where models design ligands specific to a biological target. Recent approaches have focused on using 3D structures of protein–ligand complexes or binding pockets (e.g., DiffDock [9], EquiBind [10], TargetDiff [11]) to either predict optimal docking poses for given molecules or generate novel molecules directly within binding sites. However, these models face several critical limitations. First, they assume static binding sites and overlook conformational flexibility and induced-fit effects, which are often essential for ligand potency. Second, they rely on paired protein–ligand structural data, which remains limited (fewer than 30,000 complexes in the PDBbind database [12]) and biased towards well-studied targets and chemotypes. Third, structure-based optimization can constrain chemical diversity, prioritizing docking fit over meaningful properties such as drug-likeness, pharmacokinetics, or novelty.

The success of AlphaFold2 [13] and large-scale protein language models [14, 15] has enabled accurate prediction and analysis of protein structure and function directly from amino acid sequences. These models capture rich biophysical and evolutionary information, enabling accurate inference of structural features, functional motifs, and binding potential from sequence alone. This paradigm shift opens the door to generative models that can operate at the proteome scale, even in the absence of experimentally determined 3D structures. At the same time, advances in cross-modal generative modeling, such as text-to-image diffusion systems like DALL-E 2 [16], Imagen [17] and Stable Diffusion [18], have demonstrated the power of conditioning the generative process on learned semantic embeddings.

At the same time, advances in discrete diffusion models have significantly broadened the capabilities and applications of generative AI models [19– Unlike autoregressive methods, discrete diffusion enables efficient, non-sequential sampling and better modeling of longrange dependencies – both crucial for generating chemically valid molecules with diverse structures. Recent works such as GenMol [24] and PepTune [25] have demonstrated that masked discrete diffusion models can effectively learn the syntax of SMILES representations, enabling the generation of valid and diverse compounds. Inspired by this success, we explore a related challenge in drug discovery: generating molecules conditioned on protein sequences.

We propose ProtoBind-Diff – a *protein-sequence-conditioned masked diffusion language model* for molecular generation. Our model conditions on the protein primary sequence, bypassing the need for 3D structures, and leverages pre-trained embeddings to capture functionally relevant context. By framing generation as a denoising process over the vocabulary of SMILES tokens, the model achieves high chemical diversity, structural novelty, and validity. At the same time, it preserves physicochemical property profiles similar to those of known actives while accurately conditioning on specific biological targets. The model is trained on over one million active protein–ligand pairs from BindingDB [26], a scale far exceeding what is feasible with structure-based datasets such as PDBbind.

Our results demonstrate that protein sequence alone, when paired with expressive generative models and pre-trained embeddings, can guide the creation of novel, bioactive small molecules, unlocking structure-independent ligand design across the full proteome. Altogether, we provide evidence that this structure-free approach may expand the generative drug design to tens of thousands of proteins that lack crystallographic data.

## II. RESULTS

We trained and evaluated ProtoBind-Diff, a structure-free, protein-sequence-conditioned masked diffusion model for molecular generation. Unlike structure-based generative methods that rely on resolved protein–ligand complexes, our model is trained directly on more than one million protein–ligand pairs from BindingDB, using only sequence-level data, Fig. 1**a**. Protein context is provided through ESM-2 [27] embeddings, which are integrated via cross-attention to guide the reconstruction of masked tokens in a SMILES string (see Methods VIII B for more details on the model architecture).

**FIG. 1.**
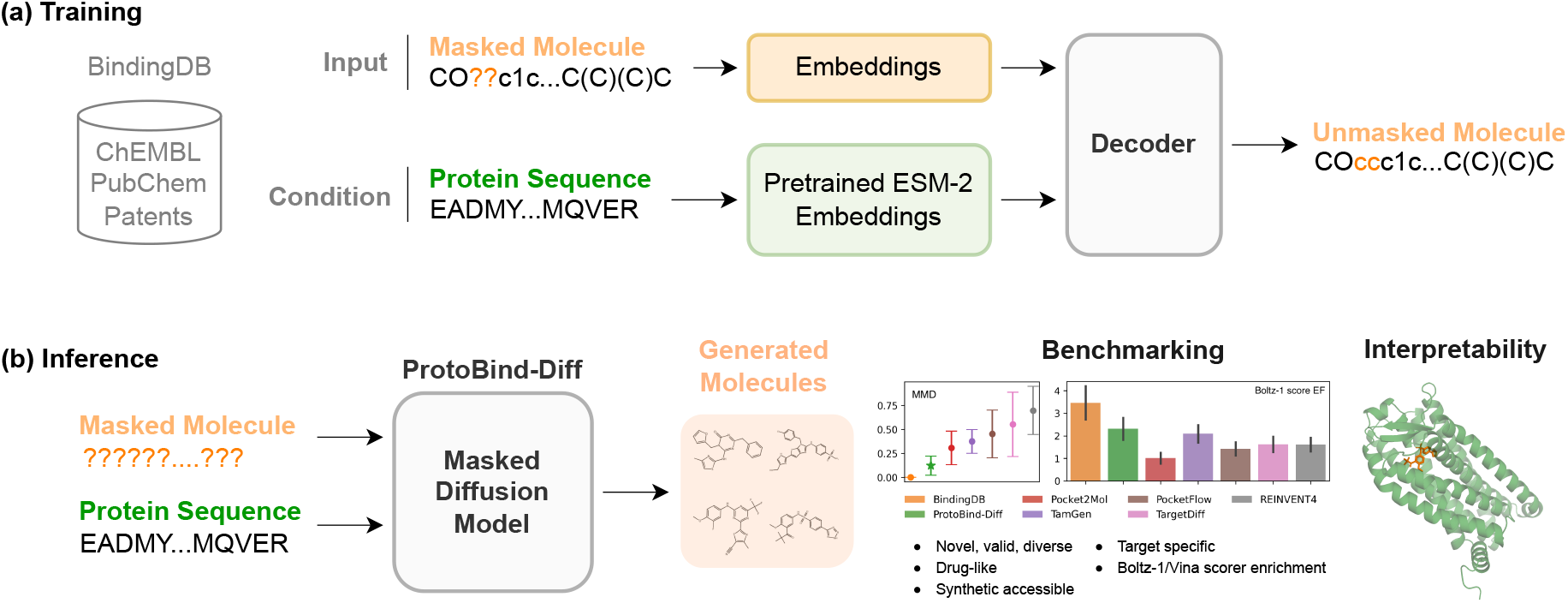
Graphical overview of ProtoBind-Diff training and application/benchmarking. **(a)** During training, ProtoBind-Diff learns to reconstruct masked molecular sequences (SMILES) conditioned on protein sequences using a denoising diffusion process. Ligands and protein targets are derived from BindingDB, with protein sequences embedded using pre-trained ESM-2 representations. **(b)** At inference time, ProtoBind-Diff generates novel, valid, and chemically diverse ligands given only a protein sequence. These ligands are compared to known actives based on key physicochemical properties, and the performance of ProtoBind-Diff is evaluated through binding affinity benchmarks against 3D structure-based generative models.

In preliminary experiments, we adapted two graphbased diffusion models, DiGress [8] and GRuM [7], to generate ligands conditioned on protein sequences, following the approach of Yang et al. [28]. While the proportion of chemically valid molecules remained high, we observed a significant drop in Tanimoto similarity between the denoised outputs and reference structures as training progressed. Furthermore, the generated molecules exhibited a strong bias toward trivial structures, often consisting of long, linear carbon chains. To address these limitations, we chose to use a text-based representation of a chemical molecule in the form of SMILES.

During inference, we generate molecules for a given protein by passing a fully masked sequence of fixed length and the target protein sequence into the trained masked diffusion model, Fig. 1**b**. The model iteratively unmasks tokens over a fixed number of denoising steps using a remasking sampler (see Methods VIII B), producing a set of candidate SMILES for each target. By tuning the remasking sampler and careful dataset curation, we achieved high chemical diversity and novelty. The generated molecules demonstrated high validity, drug-likeness, and physicochemical property distributions closely aligned with those of active molecules in BindingDB, outperforming structure-based generative baselines.

To assess target specificity, we projected the generated compounds into a UMAP space trained on active molecules and evaluated cluster separation. Binding affinity was assessed using both classical docking (AutoDock Vina [29]) and a deep learning model (Boltz-1 [30]). We further explored the interpretability of attention maps, showing emergent alignment with functional binding residues despite the absence of structural supervision.

### A. Properties of generated molecules

Following Liu et al.[31], we selected 12 diverse protein targets for the test set, spanning the 7 most common protein families according to the ChEMBL [32] protein classification. From the intersection of the CrossDocked2020 [33] and BindingDB datasets, we chose 6 ‘easy’ targets with over 1,000 training examples (ESR1, HCRTR1, JAK1, P2RX3, KDM1A, IDH1), and 6 ‘hard’ targets with few examples (RIOK1, NR4A1, GRIK1, CCR9, FTO, SPIN1). See Table S1 for annotation details.

To assess the overall performance of ProtoBind-Diff, we compared it with three recent generative models that sample molecules based on 3D protein pockets and have demonstrated strong performance [31]: PocketFlow [34], Pocket2Mol [35], and TargetDiff [11]. We also added TamGen [36] as a newer model that leverages pretrained SMILES embeddings from PubChem. In addition, we selected REINVENT4 [37], a model that generates molecules based on desired chemical properties without conditioning on protein targets. For each target and model, we generated 1,000 SMILES strings to evaluate the percentage of unique, diverse, and novel molecules.

During the validation phase, some generated SMILES were found to be invalid or duplicated, as can be seen in Fig.2**a**-**b**. These samples were excluded from further analysis. Our model demonstrates reasonable validity and uniqueness scores. We observed that validity can be improved by tuning the parameters of the remasking sampler during the generation step (see Methods VIII B); however, this comes at a trade-off against other molecular properties. For instance, TamGen produces more than 98% valid molecules, but both uniqueness and diversity are relatively low. PocketFlow achieves the best performance in terms of validity and uniqueness, but it predominantly generates molecules with lower molecular weight compared to active compounds (see Fig.2**g**), suggesting a tendency to favor simpler structures.

**FIG. 2.**
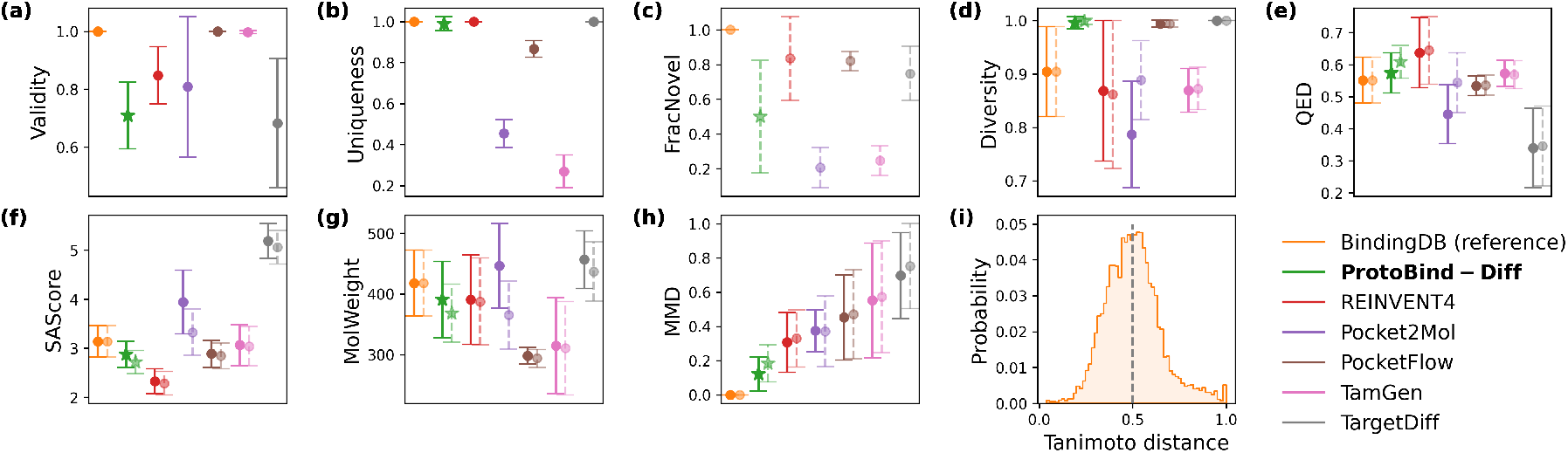
**(a)**-**(h)** The bar plots illustrate the comparison between the general chemical properties of all generated molecules (bright circles) and those that passed through the novelty filter (pale circles). The results are displayed for all generative models, with each data point representing the average performance over 12 test-set targets. All properties except validity were calculated after the standardization and duplicate removal steps. Greater similarity to the BindingDB (reference) indicates better generative quality. **(i)** The Tanimoto similarity (*T*_sim_) between molecules generated by the ProtoBind-Diff model and active molecules from BindingDB (the novelty cutoff *T*_sim_ = 0.5 is highlighted by the gray dashed line).

One of the primary objectives in drug discovery is to generate novel compounds that are structurally distinct yet retain activity against a given protein target. To assess this, we benchmarked model outputs under a molecular novelty constraint, defining novel molecules as those with a maximum Tanimoto similarity (*T*_sim_) below 0.5 to any active compound for the same target in BindingDB (Fig.2**i**). The Tanimoto similarity metric quantifies the overlap of chemical substructures encoded in bitwise vectors, where a value of 1 indicates identical compounds and 0 indicates completely dissimilar molecules. We refer to the fraction of such structurally novel molecules as the FracNovel metric (Fig. 2**c**).

Figure S2 presents Tanimoto similarity histograms across all models and targets. Interestingly, models trained on CrossDocked2020 sometimes generate compounds highly similar to known actives in BindingDB, potentially indicating overfitting. ProtoBind-Diff, Pocket2Mol and TargetDiff tend to produce molecules with high similarity to known actives for ‘easy’ targets, as indicated by a histogram peak shifted toward 1 Conversely, for ‘hard’ targets, these models generate compounds with lower similarity, shifting the histogram toward 0. We interpret high similarity as a potential sign of overfitting, while complete dissimilarity may suggest a lack of protein-specific conditioning, an issue particularly evident in models like PocketFlow and TamGen in Fig.S2. Furthermore, unconditional REINVENT4 model faces greater challenges in achieving an optimal similarity balance. REINVENT4 also tends to generate molecules that are more drug-like and easier to synthesize than reference actives, indicating a preference for chemically simpler compounds. This may reflect a bias toward general drug-likeness, placing model’s outputs further from the distribution of known actives. Finally, we observed that some models, such as Pocket2Mol, exhibit a clear gap between the chemical properties of all generated molecules and those of only novel ones, Fig.S4**d-g**.

To assess how closely the chemical property distributions of generated molecules match those of known active (reference) molecules, we computed the Maximum Mean Discrepancy (MMD) metric. MMD quantifies the difference between two distributions: a lower value indicates greater similarity, suggesting that the model replicates the property distribution of real molecules more accurately. The chemical properties used to compute MMD are detailed in Methods VIII C, and the results for all targets are shown in Fig. S3-S4. As observed, while other models sometimes outperform ProtoBind-Diff on individual descriptors, our model produces molecules whose overall chemical descriptor distributions are more closely aligned with those of the reference compounds.

### B. Structure-Based Evaluation of Generated Ligands

In the absence of experimental binding affinity data for the generated molecules, we evaluated structural plausibility using two distinct methods: classical molecular docking with AutoDock Vina and Boltz-1, a deeplearning model for biomolecular interaction prediction. Docking was performed using the standard AutoDock Vina protocol, with the binding site defined by the position of reference ligands in experimentally determined structures from the Protein Data Bank (PDB) [38] (see Table S1). All generative models performed well on targets where docking effectively distinguished active from inactive compounds — for example, ESR1, GRIK1, and CCR9 (Fig. S5). However, in most cases, docking exhibited poor discriminatory power. For example, with targets such as P2RX3, KDM1A, IDH1, RIOK1, NR4A1, FTO, and SPIN1, the difference in average docking scores between active and inactive molecules were not statistically different. Notably, for several of these targets, Pocket2Mol and REINVENT4 achieved significantly lower docking scores than all other models and even true active compounds, e.g. KDM1A, IDH1, RIOK1 and SPIN1.

For all methods, the docking scores of generated compounds varied substantially across targets. Overall docking performance is summarized in Fig. 3**a** using the enrichment factor (EF), which quantifies whether the concentration of predicted active molecules in the observed set is higher (EF > 1) or lower (EF < 1) than in the reference set. EF was computed as 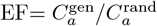, where 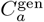 and 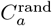 denote the fractions of active molecules (defined as having a Vina score < 10 kcal/mol) in the generated set and in the random subset of all active molecules from BindingdDB, respectively.

**FIG. 3.**
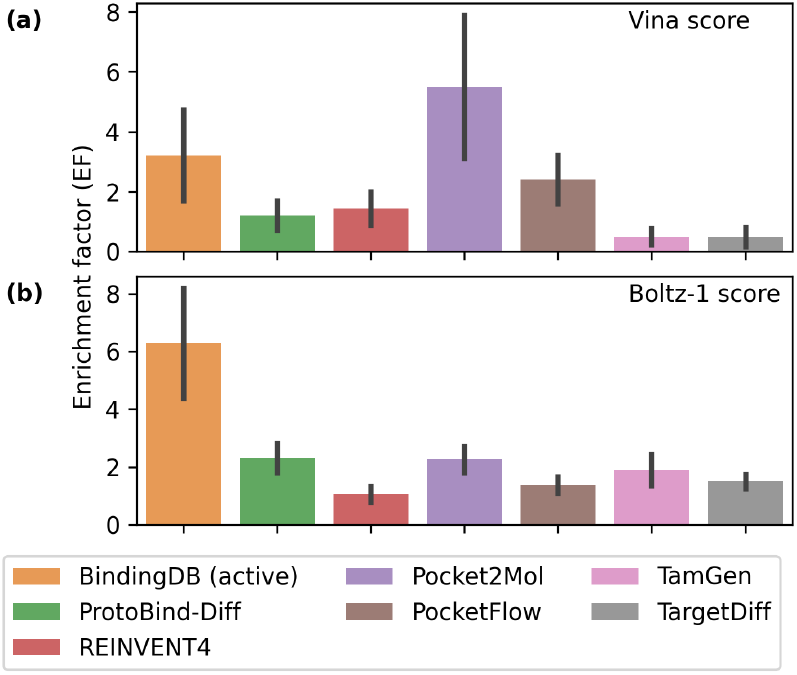
Enrichment Factor (EF) analysis of AutoDock Vina and Boltz-1 **(b)** scorers for identifying active molecules above thresholds compared to randomly selected active molecules from BindingDB. Thresholds used: AutoDock Vina docking score < –10 kcal/mol, Boltz-1 iPTM score > 0.85. Error bars represent the standard error of the mean (SEM). Data per target is presented in Tables S4-S3

Based on docking EFs, ProtoBind-Diff ranked below Pocket2Mol and PocketFlow. Notably, Pocket2Mol exhibited a surprisingly high EF, surpassing even that of the true active molecules. We attribute this to the fact that both Pocket2Mol and PocketFlow were trained on the CrossDocked2020 dataset, which, although based on crystallographic structures, was heavily augmented (by a factor of 100) with Vina-generated poses. This likely led to overfitting, causing the models to preferentially generate molecules that score well under Vina. Conversely, the relatively low EF observed for true actives suggests that the Vina scoring function may not align well with actual binding activity.

To complement docking, we applied the Boltz-1, a recent open-source deep learning model for proteinligand structure prediction, to the same sets of generated molecules and targets. Boltz-1 was used to predict ligand-protein complexes, providing an interface predicted TM-score (ipTM), a confidence metric that estimates the structural plausibility of the predicted binding interface. In our experiments, ipTM scores effectively distinguished active from inactive molecules across a wide range of targets (Fig. S6).

For nearly all ‘easy’ targets (ESR1, HCRTR1, JAK1, KDM1A, IDH1, P2RX3), ProtoBind-Diff produced ipTM score distributions that were comparable to or better than those of structure-based models, including PocketFlow, Pocket2Mol, and TargetDiff. On ‘hard’ targets, ProtoBind-Diff achieved the top or near-top ipTM scores for SPIN1, GRIK1, RIOK1, CCR9 and NR4A1.

EF results based on Boltz-1 predictions are presented in Fig. 3**b**. Compared to docking, Boltz-1 ipTM scores better discriminated actives from inactives, yielding higher EFs on BindingDB active molecules (see Tables S4 and Figure 3**b**). This suggests Boltz-1 captures more biologically meaningful interaction patterns than traditional docking algorithms. Notably, ProtoBind-Diff achieved the highest EF, closely followed by Pocket2Mol.

### C. Target specificity

The discovery of novel chemical entities requires selecting molecules that not only exhibit the desired protein activity but also possess diverse chemical scaffolds. To assess the ability of models to generate such novel molecules within a given protein class, we filtered all generated compounds to include only those with a maximum similarity below 0.5 to known actives from BindingDB, and projected them onto a UMAP plot for comparison with reference data (Fig. 4). Since structurally similar compounds often exhibit similar biological activity profiles [39], we interpret clusters in the UMAP representation as indicative of potential bioactivity similarity.

**FIG. 4.**
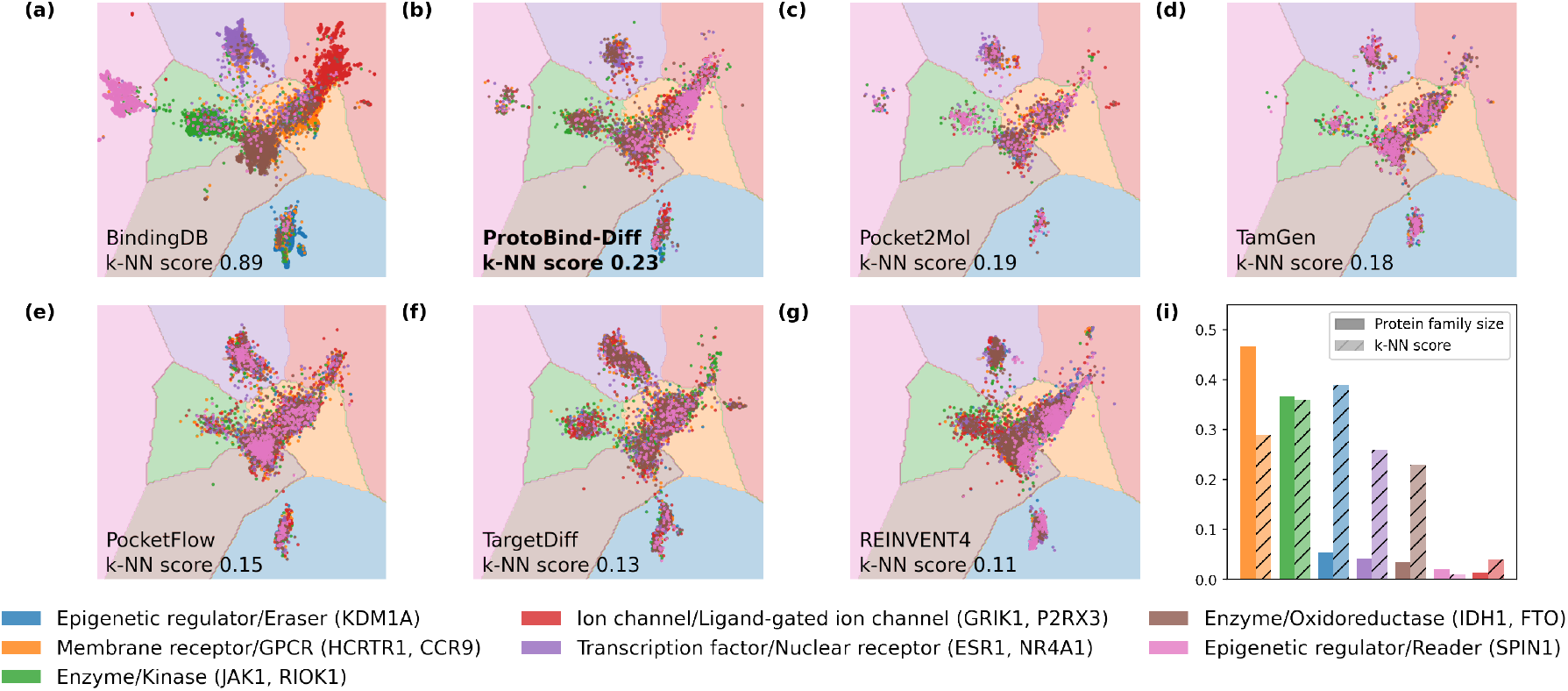
**(a)** UMAP visualization of molecular fingerprints of active molecules from the BindingDB colored by protein families according to the L1/L2 ChEMBL protein classification (bright markers, colored by protein family). The shaded areas (colored by a protein family) represent k-NN classifier decision boundaries for corresponding families trained on UMAP projection into 2D space. **(b)**-**(g)** The pre-trained UMAP transformation applied to generated novel molecules (Tanimoto similarity to known active *T*_sim_ < 0.5), which are represented by bright markers. The k-NN decision boundaries were inferred from the classifier trained on active molecules. Overall score: mean k-NN accuracy across seven protein families, where higher values indicate superior performance. **(i)** Normalized number of samples in the training set (filled boxes) and corresponding k-NN accuracy (hatched boxes) for ProtoBind-Diff model.

The UMAP model was trained on Morgan fingerprints of active molecules from BindingDB (see Methods VIII D). A k-nearest neighbors (k-NN) classifier, trained on the resulting UMAP embeddings, was used to assess the degree of separation between protein classes among all generated molecules. The classification accuracy was then used as a proxy for target-specific separability.

In the UMAP embeddings of reference actives from BindingDB (Fig. 4**a**), distinct separation by protein class is evident, with clearly defined boundaries, strong intra-class cohesion, and high k-NN classification scores. In contrast, while the embeddings of the generated molecules retain some structural similarity to the reference space, they show reduced inter-class separation. For instance, the best k-NN classification accuracy among generated molecules was only 0.23, achieved by ProtoBind-Diff (Fig. 4**b**).

Across all generative models, including ProtoBind-Diff, we observed a tendency to produce molecules resembling those of the Membrane receptor (orange area in Fig. 4) and Enzyme (brown and green areas in Fig. 4) protein classes, despite protein-specific conditioning during generation. These two classes are among the most overrepresented in public datasets, Fig. 4**i**, suggesting a generation bias toward the most frequently seen classes. On the other hand, we also obtained promising results for both highly represented classes (GPCR, Kinase) and less represented ones (Eraser, Nuclear receptor, Oxidoreductase), Fig. 4**i**. Additionally, certain protein families with low molecular diversity, such as KDM1A, allow the model to more easily generate compounds with matching structural features.

In summary, the target specificity of generated molecules is strongly influenced by the presence of similar examples in the training set, with a clear trend indicating that larger numbers of training samples lead to improved target specificity.

### D. Attention-Based Binding Site Analysis

To investigate whether ProtoBind-Diff captures interpretable patterns of protein–ligand interaction, we analyzed the attention heads in the final decoder layer using 1,843 BioLiP-2 annotated proteins [40]. Attention weights were used as predictors of binding site residues (see Methods VIII G). Figure 5**a** presents a bar plot showing the mean ROC–AUC for each attention head, with the standard errors across the annotated proteins. Attention head 8 yielded the highest ROC–AUC of 0.716 *±* 0.003. For comparison, a linear classifier trained on ESM-2 embeddings (using a similarity-based train/test split) yielded a ROC–AUC of 0.849 *±* 0.003. Since ProtoBind-Diff was not trained on BioLiP-2 residue labels or other binding pocket data, the strong performance of attention head 8 indicates that the model independently learns to focus on structurally relevant regions without explicit supervision. This suggests that the model’s attention mechanism encodes biophysically meaningful patterns.

**FIG. 5.**
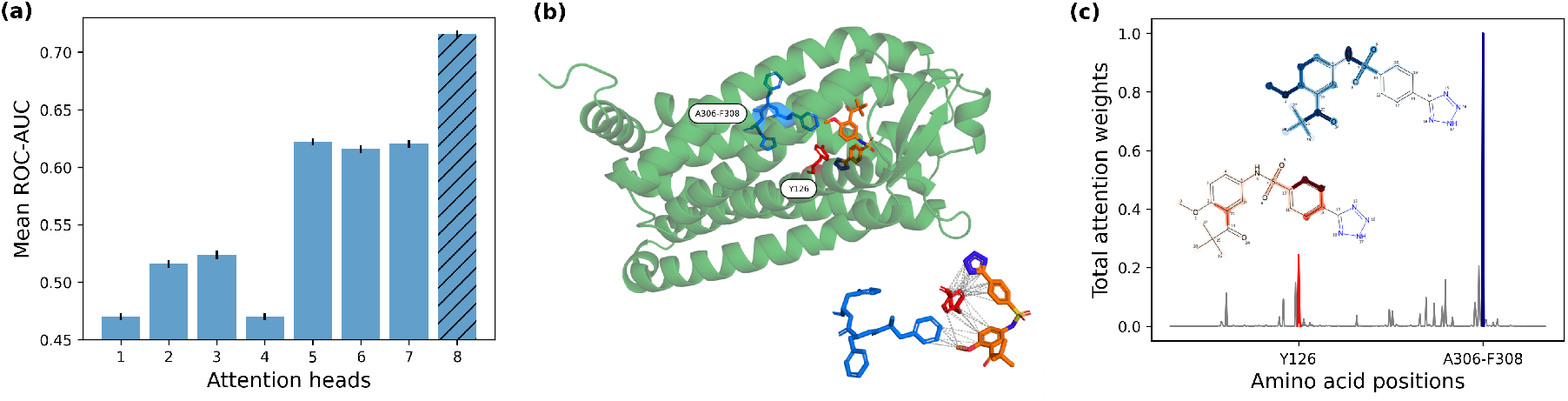
Interpretability of ProtoBind-Diff attention. **(a)** Mean ROC-AUC (± SEM) for binding-site detection for the eight attention heads of ProtoBind-Diff, averaged over 1,843 annotated sequences. Head 8 shows the highest ROC-AUC of 0.72. (b) Predicted pose of a ligand generated by ProtoBind-Diff (orange) in complex with CCR9 protein (green). The structure was generated using Boltz-1 model. Inset: Predicted binding interactions between the ligand and amino-acid contact residues, based on a 5 °A distance cutoff. **(c)** Attention weights from head 8 of ProtoBind-Diff, averaged over ligand tokens and plotted against residue positions in the protein sequence. Atoms in the molecular graph are colored with intensity proportional to their attention weights. Peaks in the amino acid sequence align with residues that are in direct contact with the ligand in the predicted pose, suggesting that the model’s attention mechanism captures spatially relevant interaction signals from sequence alone.

Figure 5 illustrates a representative case study involving the GPCR CCR9, with a ligand generated by ProtoBind-Diff that achieved a high Boltz-1 ipTM score. The protein–ligand complex was predicted using the Boltz-1 model based on the CCR9 receptor sequence from PDB entry *5LWE*. The predicted binding pose highlights specific contact residues surrounding the ligand (Fig. 5**b**).

We further extracted attention maps from attention head 8 of the final transformer block, averaging the weights across all ligand tokens to derive a per-residue weight vector.

Remarkably, the attention profile over the protein sequence (Fig. 5**c**) exhibits distinct peaks at amino acid positions that are close to the contact residues (Fig. 5**b**) and also captures ligand substructures aligned with these residues. We further quantified the contributions of individual atoms in the ligand based on their attention weight values. These results demonstrate that ProtoBind-Diff’s cross-attention mechanism effectively integrates protein sequence information and ligand structural features, aligning with biophysically meaningful interactions that mediate ligand recognition.

## III. DISCUSSION

While recent advances in 3D structure-conditioned ligand generation have shown some promise, benchmark studies reveal that these methods still suffer from issues including limited drug-likeness and synthesizability of generated molecules [31]. To address these challenges, we propose ProtoBind-Diff, a novel sequence-conditioned diffusion framework for molecular generation, which is trained entirely without 3D structural data. In doing so, it directly addresses a central bottleneck in generative drug discovery: the limited availability, bias, and rigidity of protein–ligand complex structures. While tools like AlphaFold-2 have solved the challenge of protein structure prediction, accurate information about ligand binding sites and conformations remains scarce, particularly for novel targets.

ProtoBind-Diff employs pre-trained language models that encode biochemical and evolutionary features, eliminating the need for experimental ligand-protein structures. Additionally, by tuning the model’s parameters and sampling strategies, combined with techniques such as dataset augmentation and balancing, we achieved robust molecular validity. The generated molecules exhibit high synthesizability and drug-likeness, demonstrating strong performance across our benchmarks.

In this work, we focused on 1D data representation: SMILES for ligands and FASTA sequence for proteins. An alternative approach, which typically performed well in the task of prediction, involves the use of graph neural networks (GNNs) [41, 42]. While recent advances in 2D graph-based diffusion models have shown promise for molecular generation [7, 8, 43], key limitations remain. Our empirical results show that graph-based diffusion models consistently fail to capture essential structural motifs, as evidenced by their tendency to generate unrealistic long carbon chains. This behaviour has been previously attributed to the limited expressivity of current graph neural networks [44, 45]. Even state-of-the-art hybrid models like NExT-MOL [46] have achieved better performance when incorporating 1D sequence representations.

Despite never having been trained on 3D structures, ProtoBind-Diff demonstrates spatial awareness. We discovered that, for a subset of approximately 1,800 test protein sequences, the model’s attention peaks for a single head often align with experimentally known binding sites. Notably, in cases such as the CCR9 receptor, amino acid positions with highest attention weights corresponded to functional residues inferred by Boltz-1 This level of interpretability suggests that the model does more than simply memorize training data, it captures meaningful biochemical patterns. Nonetheless, its ability to accurately associate ligand substructures with annotated binding sites remains limited and is far from perfect.

To evaluate the generated molecules in the absence of reliable experimental data, we used two benchmarks: traditional docking and Boltz-1 ipTM score. ProtoBind-Diff achieved high Boltz-1 scores on both ‘easy’ and ‘hard’ targets, suggesting that the generated molecules are likely to be active. In contrast, structure-based models trained on crystallographic and docking-generated poses (e.g., Pocket2Mol) achieve better docking scores than ProtoBind-Diff in classical docking benchmarks. However, the reliability of these docking scores is questionable, as models like Pocket2Mol may be overfitted to Vina-generated poses. This aligns with concerns raised in recent benchmark studies [31], which suggest that docking scores may be an unreliable metric for evaluating generative models.

When evaluated with Boltz-1 score, this advantage narrows substantially. The agreement between Boltz-1 and reference data with active ligands was more stable than with classical docking, which exhibited high variance. This reinforces the view that Boltz-1 score captures more biologically relevant binding compatibility.

Structural features of active molecules from BindingDB showed clear separation between protein classes, which may reflect genuine patterns of molecular recognition. However, this separation could also be an artifact of the dataset, as most actives were tested against a limited set of targets—primarily kinases and GPCRs. In contrast, generated molecules exhibited substantially weaker class distinction, with ProtoBind-Diff achieving the highest k-NN classification score of only 0.23. All models displayed a notable bias toward membrane receptors and enzymes, mirroring their overrepresentation in the training data. Despite this, we observed promising generation across both common (e.g., GPCRs, kinases) and underrepresented protein families (e.g., nuclear receptors, erasers, oxidoreductases), especially for targets with inherently low molecular diversity, such as KDM1A. These findings underscore both the limitations and the potential of current generative models, suggesting that improving scaffold diversity and class specificity may be achievable through broader and more balanced training datasets.

Beyond benchmarking, ProtoBind-Diff enables practical workflows that are otherwise inaccessible. Although structure-based models like Pocket2Mol and TargetDiff require protein–ligand complex structures for training, ProtoBind-Diff learns to generate bioactive ligands from the sequence alone, unlocking scalable ligMand discovery across the entire proteome. This capability is especially valuable for targets lacking structural data, such as orphan GPCRs, disordered proteins, fast-evolving pathogens, and neglected disease targets, areas where structure-based methods often fall short. Moreover, ProtoBind-Diff can generate candidate compounds within days, significantly accelerating the early stages of drug discovery.

These use cases illustrate the power of generative models that operate directly on protein sequences – scalable, rapid, and agnostic to structural uncertainty. Nonetheless, limitations remain. ProtoBind-Diff lacks fine-grained structural resolution and cannot yet generate 3D poses. While sequence embeddings from models like ESM-2 capture domain-level features and functional context, they omit pocket-specific details, such as local electrostatics, solvent exposure, or induced fit. We anticipate that coupling ProtoBind-Diff’s generation with structure-aware post-processing, such as conformer generation, docking, or hybrid scoring, could yield synergistic performance gains.

Together, our findings demonstrate a new type of *biological foundation models*: scalable, generalizable architectures trained on large-scale biological data that can support multiple downstream tasks. While further calibration and screening integration are needed for full deployment, this model lays a foundation for generative design at the proteome scale. We foresee its future role in multi-target hit-finding, drug discovery against neglected diseases, and pandemic response – domains where the next wave of generative biology will find its greatest impact.

## IV. ACKNOWLEDGMENTS

We thank Ilya Eder for developing the initial version of the codebase, which served as a foundation for this work. We also thank Kirill Denisov and Daniel Igumnov for carefully reading the manuscript and providing valuable feedback and suggestions.

## VI. DATA AVAILABILITY

All data used in this study are publicly available. Molecule–protein interaction data were obtained from the BindingDB database (https://www.bindingdb.org), an open-access resource for binding affinity data. No proprietary datasets were used.

## VII. CODE AVAILABILITY

The source code for this study is freely available on GitHub https://github.com/gero-science/ProtoBind-Diff.git.

## VIII. MATERIALS AND METHODS

### A. Data Preparation

We used the BindingDB database from February 2025, containing 3,010,313 measurements, of which 1,311,211 unique compounds, 9,524 unique targets from various assays and families. Ligands were presented in SMILES format and proteins were presented in the form of amino acid sequences. Before training, the data was cleaned using the following procedure:

1. Sequences lacking a UniProt ID (according to the EMBL-EBI database), with unknown organism source, or belonging to very rare clusters were removed,
2. Cytochrome P450 and Albumin were excluded from the analysis due to their non-specific binding to all ligands,
3. All invalid SMILES and SMILES containing very rare tokens (occurring fewer than 100 times in the original dataset) were removed.,
4. Only sequences with lengths between 50 and 1,500 amino acids and ligands containing between 10 and 80 atoms were retained. Very short sequences do not form stable binding pockets, while very long protein chains and large ligands were excluded to ensure the model fit into GPU memory.

For the purpose of replicating the true data distribution during training, only active instances were chosen. Binary labels were subsequently assigned based on the following criterion: a molecule was classified as active if at least one of the *K*_*i*_, *K*_*d*_ or *EC*_50_ values exhibited an activity below 1 *µ*M. The resulting dataset included 1,167,809 samples.

Protein sequences were encoded using a pre-trained protein language model. ESM-2 embeddings, characterized by 650 million parameters, 33 model layers, and a dimension of 1280, were selected for this purpose. Research indicates that increased embedding dimensionality generally corresponds to enhanced model performance. However, in order to strike a balance between quality and computational efficiency, we chose embeddings with an intermediate size. The SMILES-formatted ligand sequences were converted to tokens using the PySMILESUtils library [47].

### B. Model Construction and Training

We define a ligand as a sequence of tokens **x** from a vocabulary of size *K*, which can be described using a categorical distribution Cat (*·*; ***π***) over *K* classes (with probabilities given by ***π****∈* Δ^*K*^), where Δ^*K*^ represents the simplex over *K* categories.

The noise scheduler, *α*_*t*_ *∈* [0, 1], was defined as a monotonically decreasing function of *t*, as described by Sahoo et al. [19]. It determines the probability of retaining the original token at each step of the forward process. The marginal of **z**_*t*_ conditioned on **x** at time *t* in this case is defined as

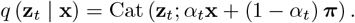

Since we are dealing with masked diffusion, ***π***can be set as ***m***and reverse process can be formulated with the following posterior:

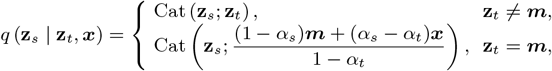

where step *s < t* and ***m***is the vector of the mask token representing a separate K-th category. It can be seen that the posterior is conditioned on **x** which is unknown, so different parametrization techniques can be used to find an approximation **x** = **x**_*θ*_(**z**_*t*_, *t*). In our case, we used the SUBS parameterization approach described in [19]. Assuming that the forward noise process is applied independently throughout the sequence, the training objective of **x**_*θ*_, approximated by the negative ELBO, is formulated as

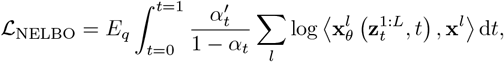

where **x**^*l*^ is a predicted value of *l*-th token, 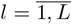.

**FIG. S1.**
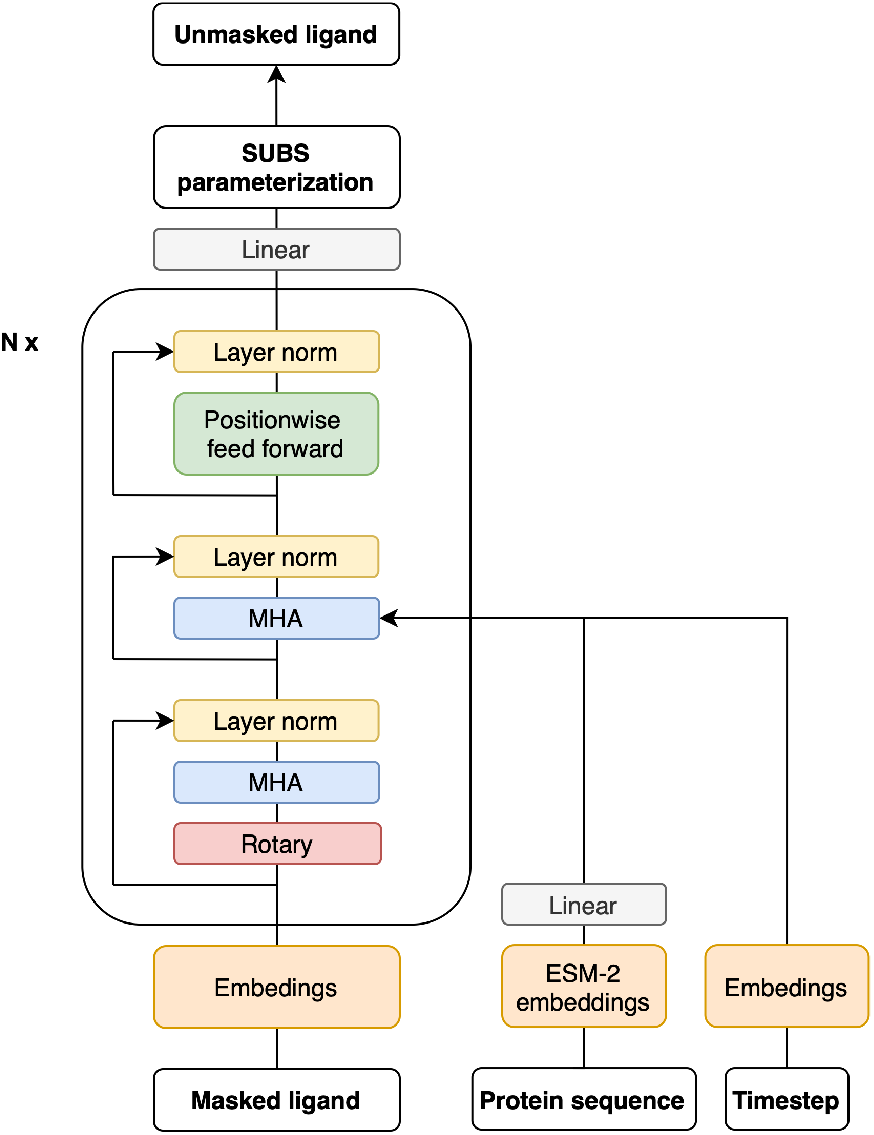
Architecture of the ProtoBind-Diff model. The masked ligand sequence is embedded, then processed through a stack of transformer decoder blocks. Each block contains rotary position embeddings[48], multi-head selfattention (MHA), cross-attention for protein sequence and timestep conditioning, followed by a normalization layer and a positionwise feedforward network. Protein sequence information is encoded using pre-trained ESM-2 embeddings, projected through a linear layer. The final output is passed through a linear layer and SUBS parameterization to predict the denoised ligand.

The model architecture consisted of several key blocks: a ligand embedding layer, responsible for converting tokens into embedding vectors; a timestep embedding module, as proposed by Peebles et al. [49]; a linear layer that transformed the dimensionality of pre-trained protein embeddings to align with the decoder’s hidden dimension; a transformer decoder which utilized rotary position embeddings [48] to reconstruct noisy ligand sequences and integrate protein sequence information via a crossattention mechanism; and finally a linear layer to adapt the decoder output to the vocabulary size (Fig. S1). We use the log-linear noise schedule proposed in [19]. By optimizing the hyperparameter space, we came to the conclusion that the best quality is achieved with learning rate 5 *×* 10^−5^, dropout 0.1, batch size 48, and the following decoder parameters: 12 layers, 8 heads and a hidden dimension 1280. The latter was chosen to match with the dimensionality of the ESM-2 embeddings. To improve generalization and reduce overfitting, we used SMILES augmentation strategy, where tokens are randomly permuted while preserving the chemical validity of SMILES as described in [50]. Furthermore, we optimised our dataset by grouping highly similar molecules from the training set within Tanimoto similarity of 0.85. Each training epoch we randomly sample one molecule from a cluster thereby reducing its effective size by approximately 2.8 times. We discovered that dataset balancing together with augmentation techniques markedly improved generalization to rare targets with limited binding data.

During the inference stage, we generated sequences of ligand molecules of a fixed length of 170 tokens by independently sampling each token. To enable the model to adjust some tokens based on their contextual relationships, we used the remasking technique introduced in [51] and nucleus sampling introduced in [52], both of which significantly reduced the number of invalid ligands generated. We experimented with all sampling approaches and parameter settings proposed in [51], and found that using nucleus sampling with a threshold of 0.9, the remdm-cap scheme with *η* = 1.0, and 250 sampling steps yielded the best results for our task.

### C. Chemical properties

To estimate the quality of generated molecules, we computed the following metrics: (1) Validity represents the fraction of valid molecules among all generated candidates; (2) Uniqueness represents the fraction of unique SMILES strings in their canonical form; (3) FracNovel represents the fraction of molecules with Tanimoto similarity less than 0.5 to the reference molecules; (4) Diversity is the number of unique clusters using Taylor-Butina [53] clustering algorithm with Tanimoto similarity cutoff 0.2 divided by the total number of samples. In addition, we computed the following molecular descriptors: (5) Molecular weight; (6) LogP (octanol-water partition coefficient); (7) Number of rotatable bonds; (8) TPSA (topological polar surface area); (9) Number of rings; (10) QED (quantitative estimate of drug-likeness) [54]; (11) SAScore (synthetic accessibility score) [55]; (12) Number of heavy (non-hydrogen) atoms; (13) Number of aromatic rings; (14) CSP3 (fraction of sp3-hybridized carbons) [56]. Descriptors (5)–(14) were used to compute the Maximum Mean Discrepancy (MMD) between the generated and reference sets. The closer these distributions are to those of known actives, the better the generation quality. All metrics and descriptors—except validity—were computed after standardization and duplicate removal, using the open-source cheminformatics library RDKit (https://www.rdkit.org).

### D. UMAP

To increase diversity, for all protein classes (ChEMBL, L2) present in the test set, active molecules were selected from BindingDB with the following criteria: (1) rare targets with a number of samples less than 2000 according to the protein families (Pfam [57]) annotation or with number of samples less than 1000 according to the ChEMBL protein classifications at level 2 were excluded; (2) unclassified proteins were removed; (3) all organisms except for eukaryotes were excluded; and (4) similar molecules (*T*_sim_ > 0.85) were removed.

Subsequently, 3,000 samples were randomly selected from each protein class in the test set. The subset was then supplemented to ensure that each target in the test set included at least 500 active molecules. For the selected molecules, we used RDKit to compute Morgan fingerprints of size 512 with radius 2. A UMAP model was then trained on these fingerprints using the following parameters: number of neighbors 50, minimal distance 0.2, spread 0.75, metric Jaccard, number of epochs 500, target number of neighbors 100, and repulsion strength 0.02. The trained UMAP was used to reduce the dimensionality of molecules’ fingerprints from all generative models for further visualization and k-NN classifier training and evaluation. The k-nearest neighbors (k-NN) classifier’s parameter for the number of neighbors was set to *k* = 50, as this configuration demonstrated optimal classification accuracy for the reference molecules.

### E. Docking Evaluation

For each benchmark target and generative model, we generated 1,000 molecules and applied a consistent selection protocol to obtain the final set. After filtering for valid molecules, we computed the Tanimoto similarity between each generated compound and all known actives for the same target. Only molecules with a maximum similarity below 0.5 were retained, ensuring structural novelty with respect to known ligands. From this subset, we randomly sampled up to 100 unique molecules per target. We used DockStream [58], a molecular docking wrapper for Python, for the automated preparation of targets, ligand embedding, and docking. The processed crystal structures and the reference ligand for pocket detection were taken from CrossDocked2020 [33] dataset. The PDB accession codes for each test target are presented in Table S1. The grid box size was 20 × 20 × 20 °A centered on the position of the center of mass of the reference ligand. The docking scores of the generated molecules were obtained for each target using AutoDock Vina version 1.2.5 [29] with default parameters unless otherwise specified.

**FIG. S2.**
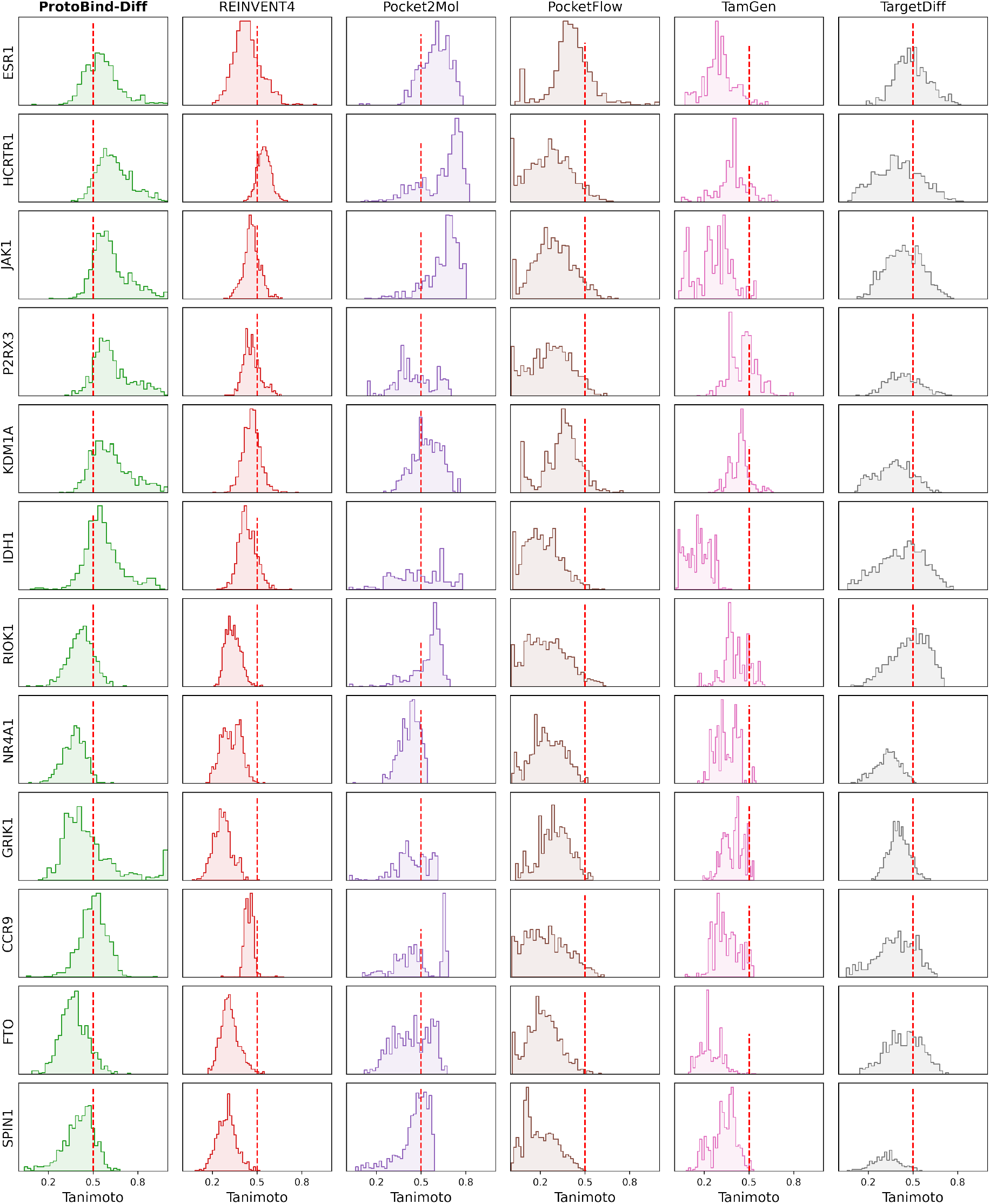
Maximum Tanimoto similarity between molecules generated by different models across 12 benchmark protein targets and known active molecules from BindingDB. Red dashed line indicates the novelty threshold (*T*_sim_ = 0.5).

**FIG. S3.**
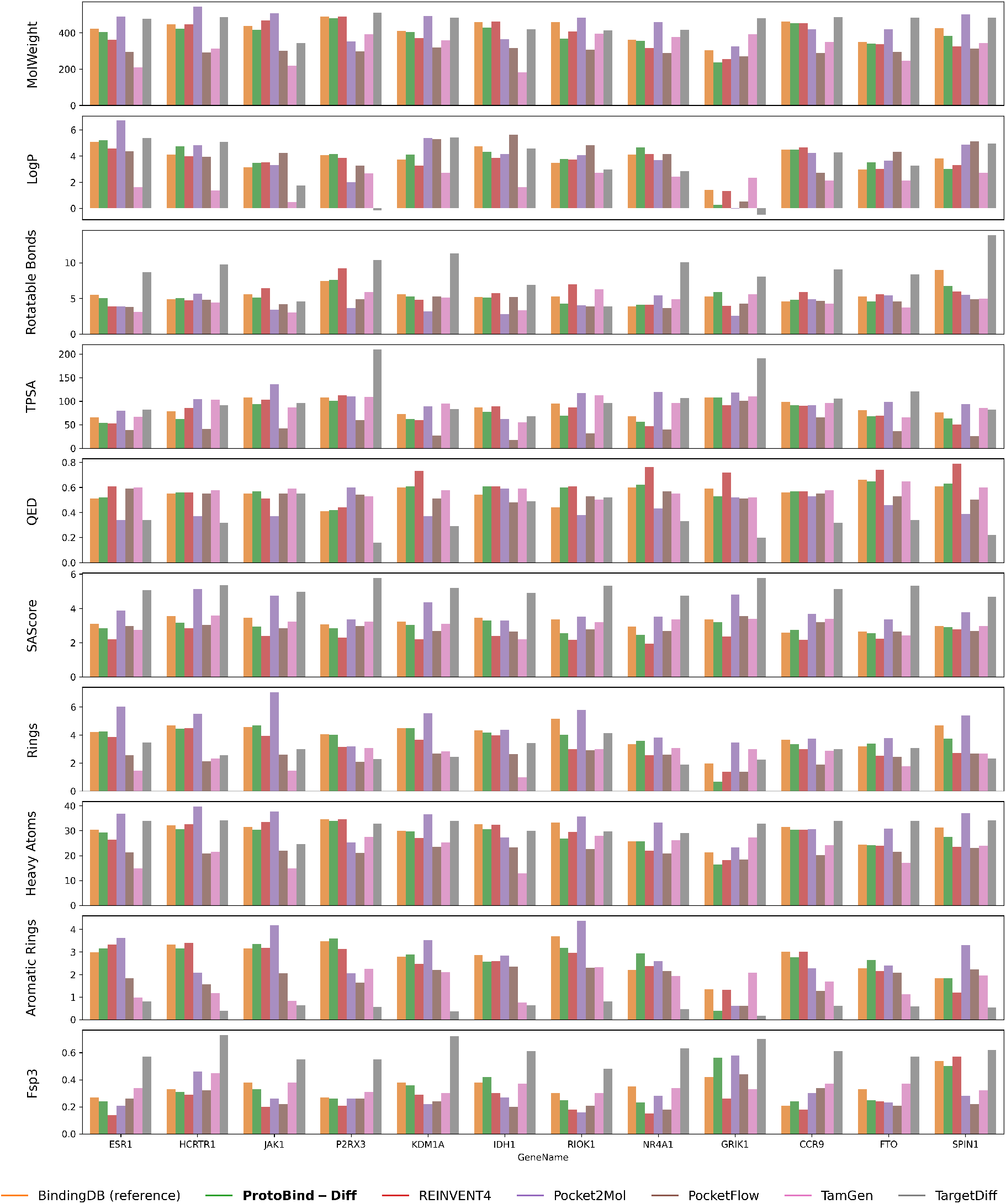
The main chemical properties of generated molecules for all targets separately from the test dataset grouped by generative models.

**FIG. S4.**
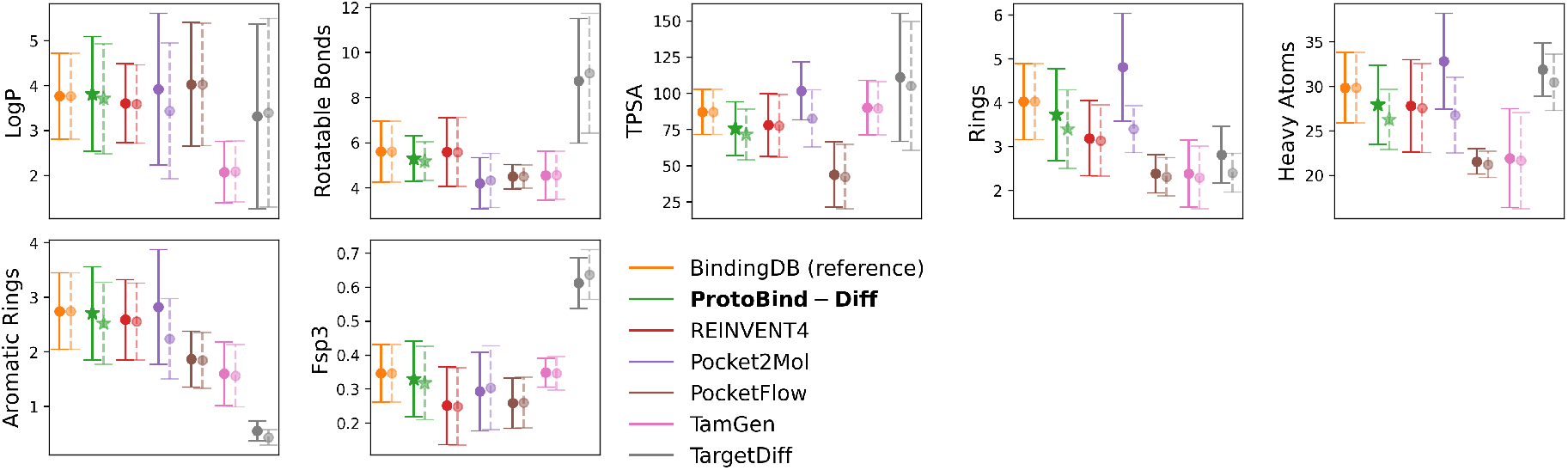
The primary physicochemical characteristics of all generated compounds (the bright circles), including those selected based on their similarity to bioactive molecules in BindingDB with a threshold of at least 0.5 (the pale circles), aggregated across all targets in the test dataset.

### F. Boltz-1 Evaluation

Boltz scores were calculated using the publicly available Boltz-1 pre-trained model, which integrates ligand preparation, pose generation, and scoring into a reproducible workflow. Boltz-1 was trained on all PDB protein–ligand complexes released before September 30, 2021, with a resolution of at least 9 °A, as described by Wohlwend et al [30]. Protein sequences from the PDB were utilized as inputs. Each ligand was scored based on the interface TM-score (ipTM) for ligand-protein complex, which estimates the confidence of interfacial structural alignment. We observed that active molecules consistently show higher ipTM scores than inactive ones (see Fig. S6), indicating that the Boltz-1 interface score can discriminate binders from non-binders. Statistical comparisons and molecules preprocessing were carried out in an analogous manner to that employed for docking protocols.

### G. Attention Visualization and Docking Analysis

From the set of canonical protein sequences of the training set, we selected those that have binding site annotations in BioLiP-2 [40], which resulted in 1843 sequences. For each selected sequence, we passed an active molecule through the model and extracted the attention weights of the final layer of the decoder. For every generated ligand, attention scores were averaged over ligand tokens to obtain a per-residue weight vector. Canonical sequences were segmented into non-overlapping 3-residue windows; each window was assigned the maximum weight among its residues. This step allows us to treat near-misses as successful binding site detections. A window was labeled ‘positive’ if any of its residues overlapped a BioLiP-annotated binding site. We then computed per-protein ROC–AUC from the window-level labels and scores. The resulting mean ROC–AUCs across all selected proteins are reported in Fig. 5**a**.

To compare our results to a baseline, we trained a simple logistic regression on ESM-2 embeddings, using a train–test split based on protein sequence dissimilarity. Protein sequences were clustered using CD-HIT [59] at 60 % identity, and clusters were randomly divided into train and test sets. After training the model to predict binding site residues, we evaluated its performance on the test set using ROC–AUC, following the same windowing procedure.

To interpret the attention weights of our model, we sampled molecules for the C-C chemokine receptor type 9 (CCR9) protein. Subsequently, a representative molecule with a high Boltz-1 ipTM score for ligand-protein complex was selected.

We focused on attention head 8 since it showed the best ROC-AUC values across all the analyzed heads. We identified peaks in the resulting profile and highlighted the corresponding values on the molecule using RDKit; these were then compared to the residues in contact with the ligand’s docked pose, which was visualized using PyMOL. [60].

**TABLE S1.**
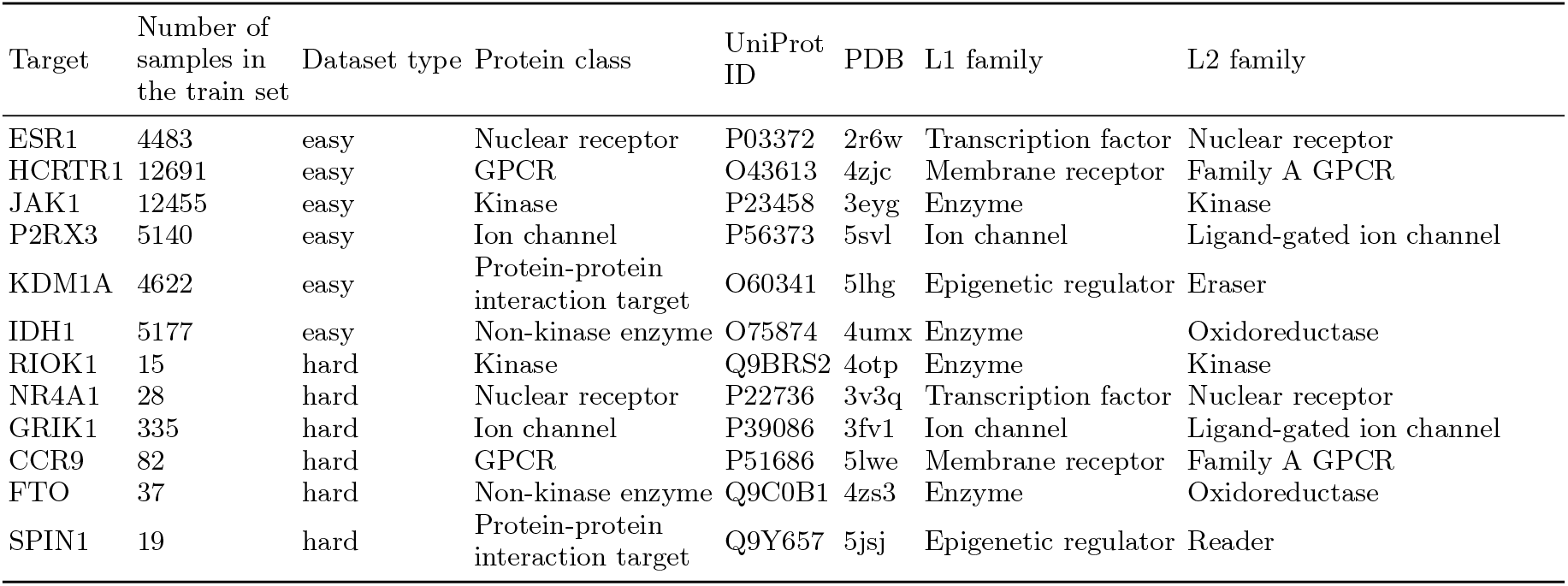
Table with annotation of chosen proteins for the test set. For each target, we listed the number of training samples available, the type of dataset (easy/hard), the protein family name, the UniProt identifier, the PDB code, and the first- and second-level family names (L1 and L2) according to the ChEMBL classification.

**TABLE S2.** Computed k-NN scores of Morgan fingerprints projected by pre-trained UMAP for all generated molecules. All family names are listed according to the L1/L2 ChEMBL classification.

| L1/L2 family name | BindingDB | ProtoBind-Diff | Pocket2Mol | TamGen | PocketFlow | TargetDiff | REINVENT4 |
| --- | --- | --- | --- | --- | --- | --- | --- |
| Epigenetic regulator/Eraser | 0.94 | **0.39** | 0.04 | 0.18 | 0.04 | 0.07 | 0.13 |
| Membrane receptor/GPCR | 0.81 | 0.29 | 0.26 | **0.38** | 0.23 | 0.28 | 0.10 |
| Enzyme/Kinase | 0.85 | **0.36** | 0.20 | 0.11 | 0.03 | 0.11 | 0.09 |
| Ion channel/Ligand-gated ion channel | 0.91 | 0.04 | **0.06** | 0.01 | 0.04 | 0.01 | 0.01 |
| Transcription factor/Nuclear receptor | 0.89 | 0.26 | **0.43** | 0.12 | 0.16 | 0.30 | 0.04 |
| Enzyme/Oxidoreductase | 0.89 | 0.23 | 0.31 | 0.44 | **0.54** | 0.14 | 0.38 |
| Epigenetic regulator/Reader | 0.96 | 0.01 | **0.03** | 0.00 | 0.00 | 0.00 | 0.00 |
| Averaged score | 0.89 | **0.23** | 0.19 | 0.18 | 0.15 | 0.13 | 0.11 |

**TABLE S3.** Enrichment factors for each target based on results of docking. The last column indicates whether the observed enrichment is statistically significant.

|  | BindingDB (active) | ProtoBind-Diff | REINVENT4 | Pocket2Mol | PocketFlow | TamGen | TargetDiff | Significance |
| --- | --- | --- | --- | --- | --- | --- | --- | --- |
| ESR1 | 18.43 | 4.95 | 4.85 | 25.37 | 7.84 | 0.00 | 3.96 | 1 |
| HCRTR1 | 1.73 | 1.21 | 2.07 | 1.23 | 0.46 | 0.04 | 0.27 | 1 |
| JAK1 | 2.12 | 0.39 | 0.96 | 0.00 | 2.80 | 0.00 | 0.17 | 1 |
| P2RX3 | 0.50 | 0.00 | 0.00 | 0.00 | 0.00 | 0.73 | 0.00 | 0 |
| KDM1A | 0.56 | 0.14 | 0.14 | 3.33 | 0.28 | 0.00 | 0.14 | 0 |
| IDH1 | 1.13 | 0.93 | 3.10 | 1.40 | 3.30 | 0.00 | 0.70 | 0 |
| RIOK1 | 6.93 | 2.91 | 0.97 | 19.16 | 7.00 | 2.91 | 0.00 | 0 |
| NR4A1 | 0.00 | 0.00 | 0.00 | 0.00 | 0.00 | 0.00 | 0.00 | 0 |
| GRIK1 | 1.04 | 0.00 | 0.00 | 1.02 | 0.00 | 0.00 | 0.00 | 1 |
| CCR9 | 5.53 | 3.76 | 5.17 | 4.93 | 1.18 | 1.88 | 0.48 | 1 |
| FTO | 0.52 | 0.16 | 0.00 | 1.29 | 1.14 | 0.00 | 0.00 | 0 |
| SPIN1 | 0.00 | 0.00 | 0.00 | 8.23 | 4.85 | 0.39 | 0.20 | 0 |

**FIG. S5.**
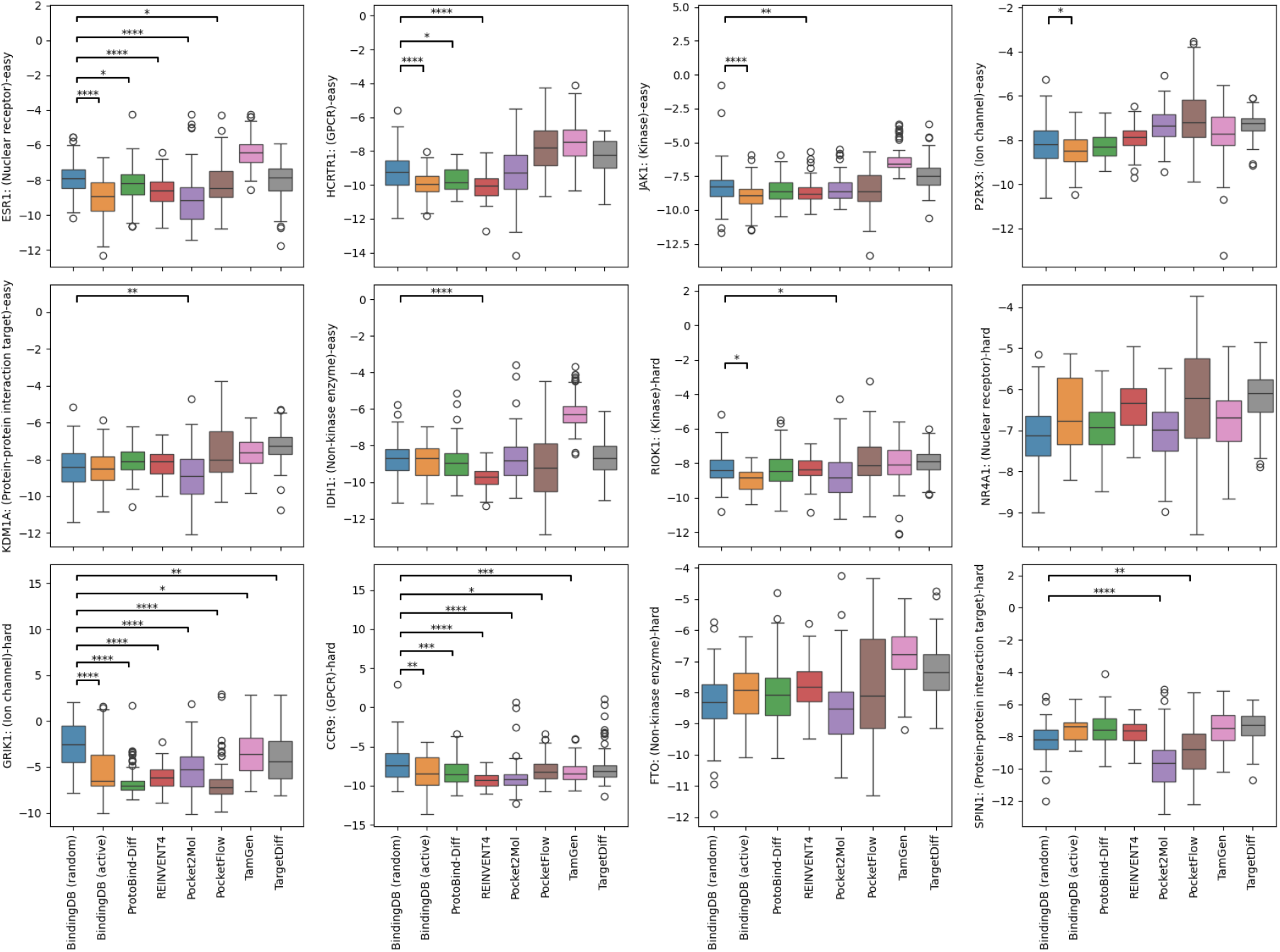
Docking scores of molecules generated by different models for 12 benchmark protein targets. Each boxplot shows the distribution of docking scores (lower is better). Statistical differences between selected model pairs were tested using the two-sided Mann–Whitney U test. Significance thresholds for adjusted p-values (Bonferroni correction): p < 0.05 (), < 0.01 (), < 0.001 (), < 0.0001 (****).

**TABLE S4.**
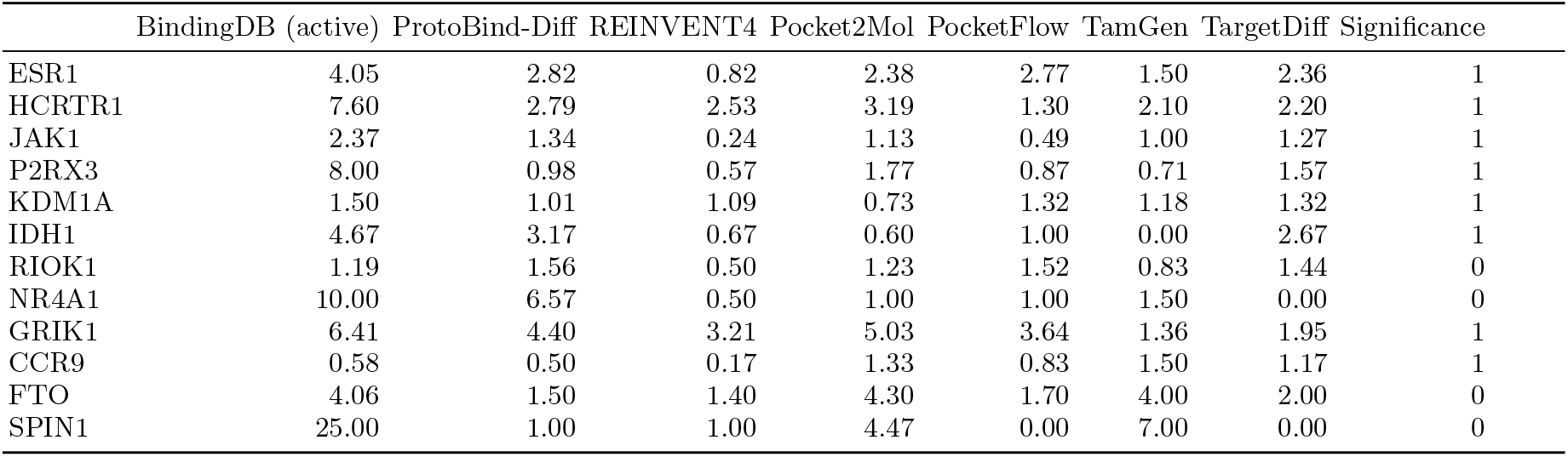
Enrichment factors for each target based on Boltz-1 ipTM scores. The last column indicates whether the observed enrichment is statistically significant.

|  | BindingDB (active) | ProtoBind-Diff | REINVENT4 | Pocket2Mol | PocketFlow | TamGen | TargetDiff | Significance |
| --- | --- | --- | --- | --- | --- | --- | --- | --- |
| ESR1 | 4.05 | 2.82 | 0.82 | 2.38 | 2.77 | 1.50 | 2.36 | 1 |
| HCRTR1 | 7.60 | 2.79 | 2.53 | 3.19 | 1.30 | 2.10 | 2.20 | 1 |
| JAK1 | 2.37 | 1.34 | 0.24 | 1.13 | 0.49 | 1.00 | 1.27 | 1 |
| P2RX3 | 8.00 | 0.98 | 0.57 | 1.77 | 0.87 | 0.71 | 1.57 | 1 |
| KDM1A | 1.50 | 1.01 | 1.09 | 0.73 | 1.32 | 1.18 | 1.32 | 1 |
| IDH1 | 4.67 | 3.17 | 0.67 | 0.60 | 1.00 | 0.00 | 2.67 | 1 |
| RIOK1 | 1.19 | 1.56 | 0.50 | 1.23 | 1.52 | 0.83 | 1.44 | 0 |
| NR4A1 | 10.00 | 6.57 | 0.50 | 1.00 | 1.00 | 1.50 | 0.00 | 0 |
| GRIK1 | 6.41 | 4.40 | 3.21 | 5.03 | 3.64 | 1.36 | 1.95 | 1 |
| CCR9 | 0.58 | 0.50 | 0.17 | 1.33 | 0.83 | 1.50 | 1.17 | 1 |
| FTO | 4.06 | 1.50 | 1.40 | 4.30 | 1.70 | 4.00 | 2.00 | 0 |
| SPIN1 | 25.00 | 1.00 | 1.00 | 4.47 | 0.00 | 7.00 | 0.00 | 0 |

**FIG. S6.**
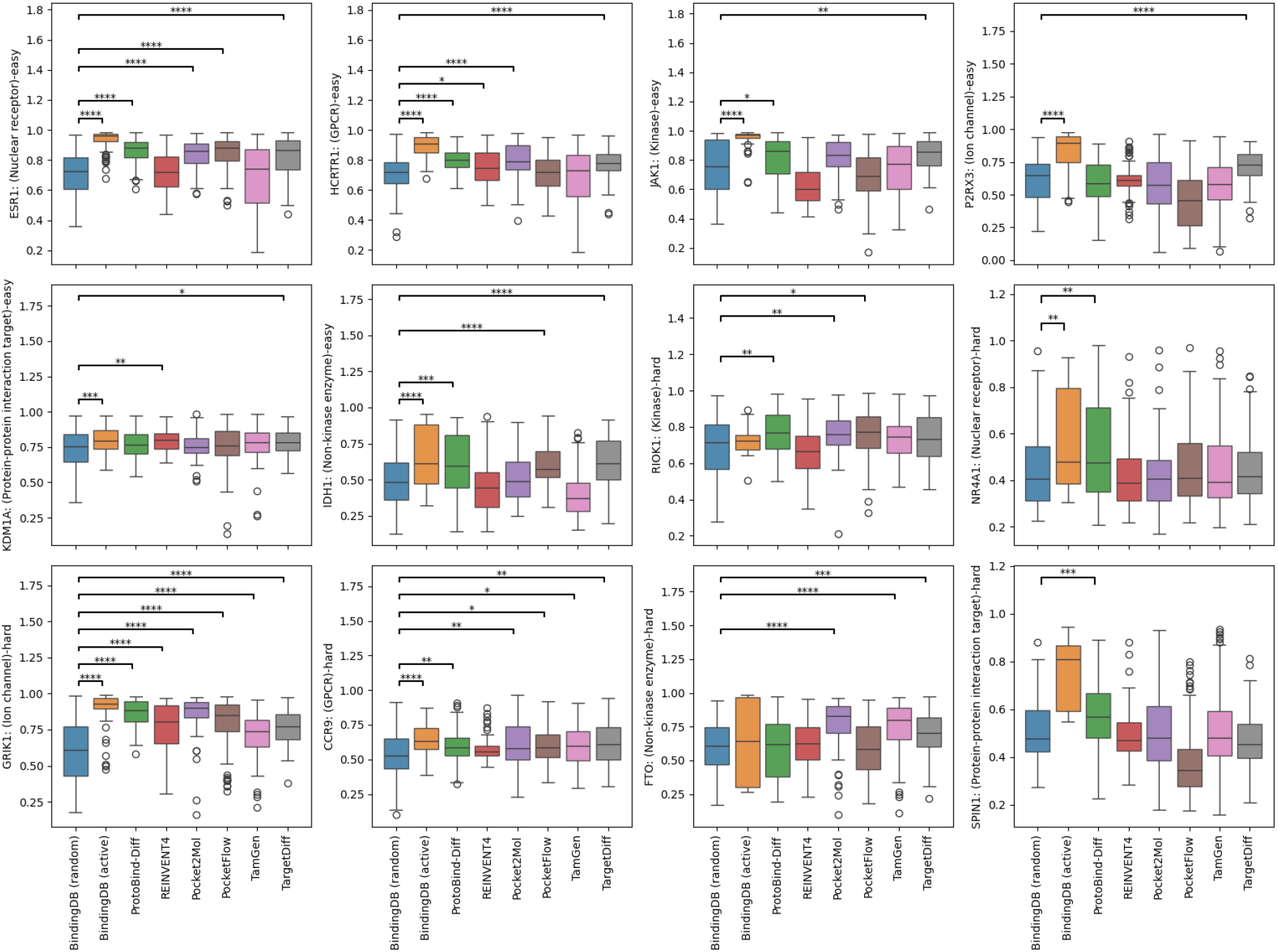
Boltz-1 scores of molecules generated by different models across 12 benchmark protein targets. Each boxplot shows the distribution of Boltz-1 scores for generated ligands targeting a specific protein, grouped by generative model. Targets are categorized as ‘easy’ (top two rows) or ‘hard’ (bottom two rows) based on training set coverage. Statistical comparisons were performed using two-sided Mann–Whitney U tests. Significance thresholds for adjusted p-values (Bonferroni correction): p < 0.05 (*), < 0.01 (**), < 0.001 (***), < 0.0001 (****).

